# Differential scanning fluorimetry as a measure of functionality of refolded anti-malarial antibody fragments

**DOI:** 10.1101/2022.09.01.504486

**Authors:** Ashwathi Valiyaparambil, Deepak K Jagannath, Vysakh K Viswanath, Naveen Kumar, Jay Prakash Shukla, Sabyasachi Pradhan, Anirudha Lakshminarasimhan

## Abstract

Transmission blocking monoclonal antibodies, 4B7 and 1245, targeting Pfs25, have been demonstrated to block the developmental stages of the malarial parasite inside the mosquitoes. In this work, anti-*Plasmodium* antibody fragments of 4B7 and 1245, designated as diabodies, were expressed in bacteria and refolded using a standardized method, with a yield of 0.1 and 0.4 mg per litre of culture, respectively. Dimeric species was detected for 4B7-Db, but not for 1245-Db, using glutaraldehyde cross-linking experiment. The quality of the purified refolded fraction was assessed with differential scanning fluorimetry (DSF). Refolded 4B7-Db, with a melting temperature of 59°C, recognized Pfs25 expressed on the surface of ookinete and zygote stages of *Plasmodium* in an immunofluorescence-based assay, whereas, 1245-Db, exhibiting a skewed melt profile, showed weak recognition. Further, refolded 4B7-Db recognized the linear epitope, on purified Pfs25 protein, both in the denatured and native state. Differential scanning fluorimetry can be potentially employed as a qualitative measure of functionality, to evaluate refolded proteins, with applicability for antibody engineering in passive immunization.

## Introduction

The significance of antibodies in therapeutics, diagnostics and imaging is exemplified by the fact that more than one hundred antibodies have been approved by the U.S. Food and drug administration (FDA) (1). From OKT3, the first monoclonal antibody (mAb) approved in 1986 (2), to dostarlimab, the hundredth in 2020 (3), the field of antibodies has been ever expanding with time. Antibody fragments granted with regulatory approvals include Brolucizumab, a Humanized scFv targeted against VEGF-A for Macular degeneration (4), Moxetumomab pasudotox, a murine IgG1 dsFv targeting CD22 for hairy cell leukemia (5) and Blinatumomab, a Murine bispecific tandem scFv targeting CD19 and CD3 for acute lymphoblastic leukemia (6). In addition to this, approval of Caplacizumab, the first nanobody in 2019 for the treatment of acquired thrombotic thrombocytopenic purpura (aTTP), has encouraged the development of antibody fragments as therapeutic agents (7). Despite their advantages like smaller size and better tissue penetration, only a handful of antibody fragments have been approved, as fragments tend to have a shorter half-life and poorer potency than their full-length counterparts. These gaps need to be carefully assessed in different therapeutic areas to identify niche areas, where in, antibody fragment-based therapeutics can be used for intervention. Antibody fragments can be engineered for improved stability and reduced immunogenicity: Moxetumomab pasudotox has a disulphide bond engineered between the Variable heavy and light chains and is conjugated to an immunotoxin (5; Fig 1A), whereas Blinatumomab is designed to be a bispecific antibody (6; Fig 1B).

**Figure 1:**
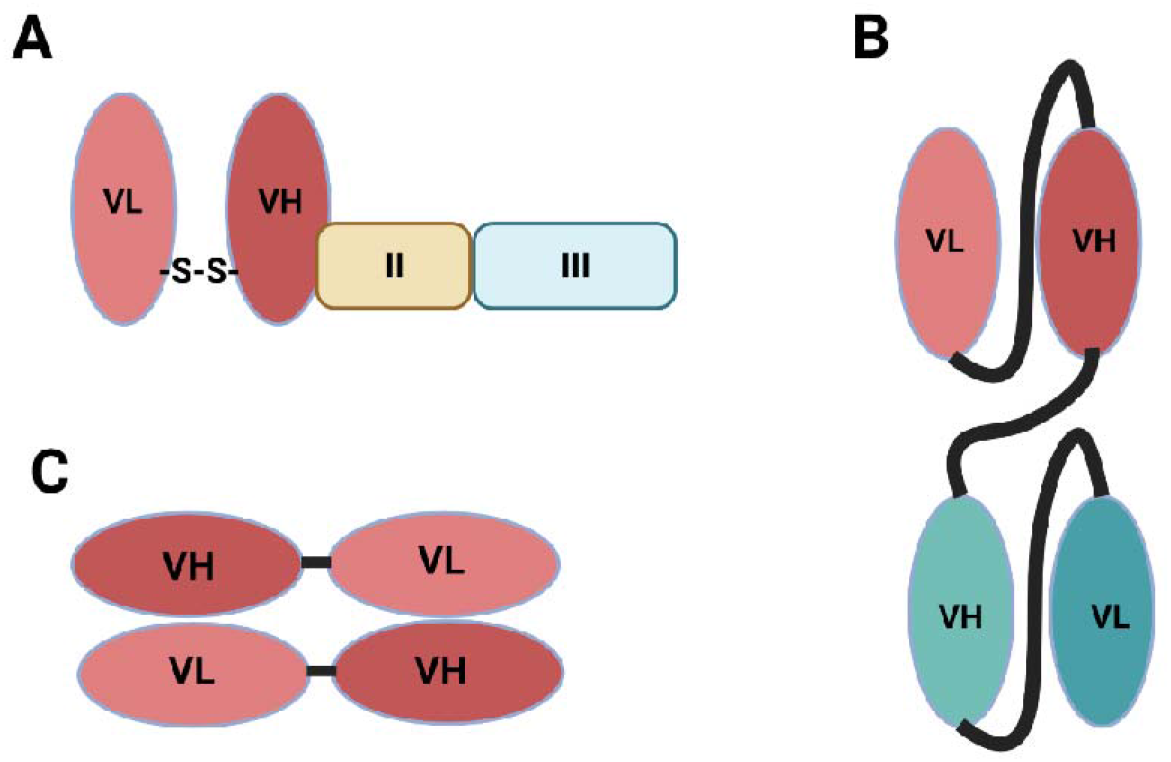
Schematic representation of engineered single chain antibody fragments. A. Moxetumomab pasudotox, an FDA approved anti-CD22 dsFv, contains an engineered disulphide bond between Position 44 of V_H_ and position 100 of V_L_, stabilizing the antibody molecule. Further, the fragment is fused to domains II and III of Pseudomonas exotoxin. B. Blinatumomab, an FDA approved anti-CD19 and -CD3 scFv, has two paratopes, each constructed by a V_H_ and a V_L_ with long linkers, conferring bi-specificity. C. Diabody is a bivalent antibody fragment, in which V_H_ from one polypeptide interacts with V_L_ from the second polypeptide, a consequence of shortened inter-variable chain linker.

Another design that has been explored is the diabody format (8-11; Fig 1C), in which a five amino acid linker (G_4_S) between the variable heavy (V_H_) and variable light domain (V_L_), prevents intrachain and promotes interchain interaction between the V_H_ and V_L_ domains (8). Despite extensive efforts put forth for the development of malaria vaccines, RTS,S/AS01, is the only approved vaccine available (12). Due the modest response observed for RTS,S/AS01, there is a need employ new interventions to combat morbidity and mortality in malaria. Clinical trials are ongoing to evaluate the protective effects of the mAb, CIS43LS, against malaria (13). Further studies are underway to engineer CIS43LS to enhance its durability (14). 4B7 (15) and 1245 (16), are transmission blocking mAbs targeting Pfs25, an extensively explored vaccine candidate for malaria (17, 18).

Prokaryotic expression systems have been used for production and characterization of antibody fragments, to screen and identify designs, exhibiting desired properties. In this study, we designed diabody (Db) counterparts of mAbs - 4B7 and 1245, recombinantly expressed them in bacteria and refolded from inclusion bodies. The quality of protein melting profiles for 4B7-Db and 1245-Db correlated with the antibody fragment’s ability to recognize surface expressed Pfs25, i.e., their functionality.

## Results

### 4B7-Db and 1245-Db were refolded using a protocol established previously for 1269-Db

pET21a vector with codon optimized DNA sequence for the cassette V_H_-G_4_S-V_L_-His_6_ from 4B7 and 1245 was expressed in *E. coli* BL21(DE3). The method optimised earlier for refolding 1269-Db (19), resulted in the recovery of the antibody fragments in solution after solubilization, refolding, dialysis and purification (Fig. 2A & 2B). After the last step (buffer exchange with G25 column), the yield of 4B7-Db and 1245-Db was found to be 0.1 and 0.4 mg/litre of bacterial culture, respectively. The *Plasmodium* antigen, MBP-Pfs25 was expressed in the periplasm of *E. coli* BL21(DE3) and purified with amylose resin and size exclusion chromatography (Fig. 2C). The monomeric fraction was purified as indicated in the previous report (19).

**Fig.2.**
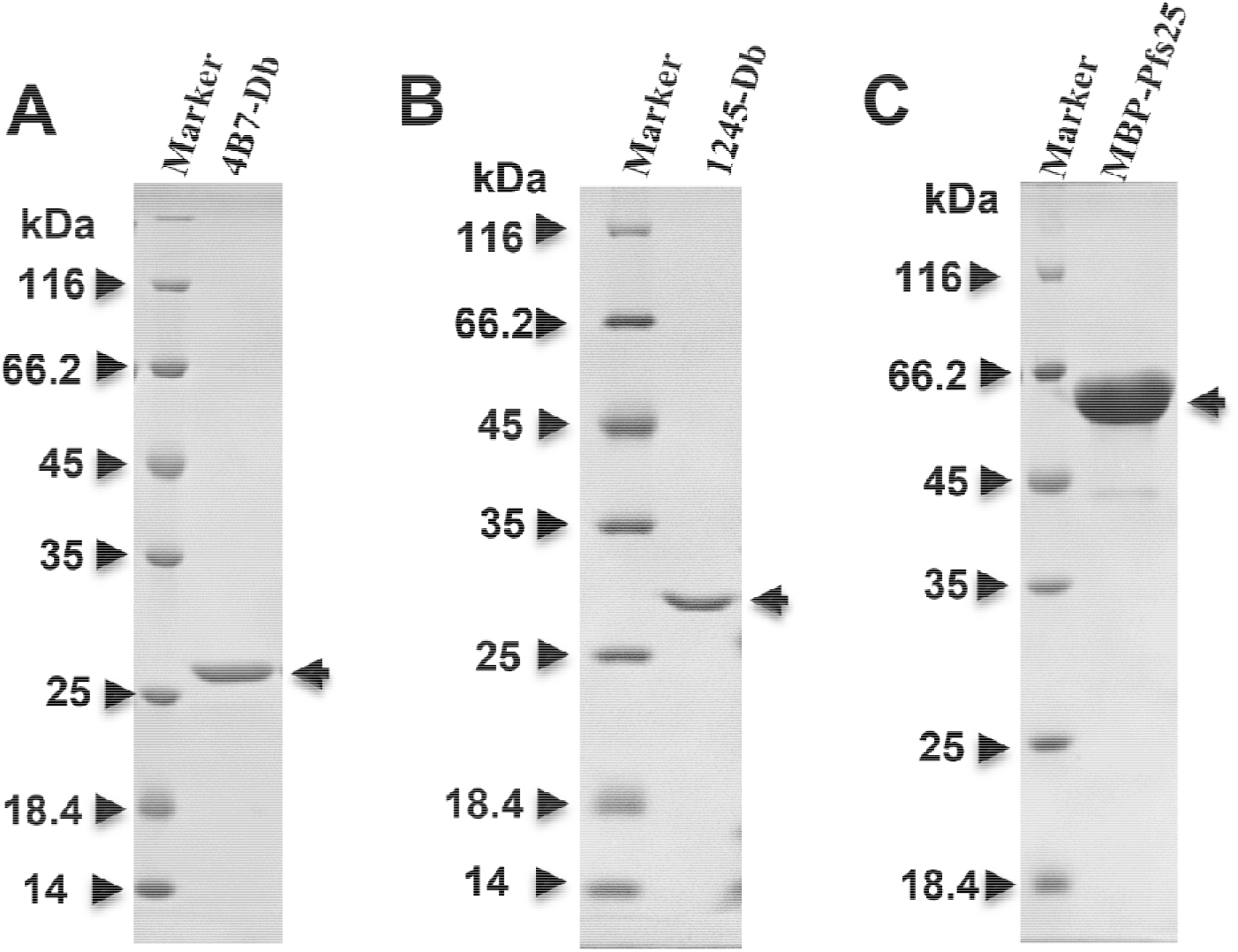
Purified antibody fragments and its concomitant antigen. Coomassie stained SDS-PAGE gel images of A. 4B7-Db, B. 1245-Db and C. MBP-Pfs25. Antibody fragments, 4B7-Db and 1245-Db were refolded from inclusion bodies, purified with affinity chromatography and G25 column chromatography (A & B), with an expected molecular weight of 26.3 and 27.9 kDa, respectively. Periplasmic expressed MBP-Pfs25 purified with affinity chromatography (C), with a molecular weight of ∼ 63 kDa. The purified antibody fragments and MBP-Pfs25 were probed with anti-His and anti-MBP antibodies.

### Refolded 4B7-Db, but not 1245-Db exhibited a sigmoidal melting profile

Refolded 4B7-Db showed an inflection temperature of 59.07 ± 1°C (n=3), which was comparable to T_i_ of refolded 1269-Db, reported earlier as 55.17 ± 0.48 °C (Fig. 3A) (19). On the other hand, high variability was observed for 1245-Db, in the replicates measured. Δ ratio of ∼0.05 along with the absence of a signature melt profile, indicated that the proportion of unfolded species in the preparation of 1245-Db was high. Although, 1269-Db, 4B7-Db and 1245-Db were refolded using the same optimized refolding condition, 1245-Db did not exhibit a standard sigmoidal profile (Fig. 3B). The amino acid sequence identities between 4B7-Db and 1269-Db, 4B7-Db and 1245-Db, and between 1269-Db and 1245-Db is 62%, 50%, and 56%, respectively. There are four positions with conserved tryptophan, in all the three antibody fragments. 1245-Db contains two additional tryptophan in the primary sequence. Additionally, 1245-Db has an alkaline theoretical pI of 8.94, as compared to the neutral to acidic theoretical pIs of 7.0 and 6.4 for 1269-Db and 4B7-db, respectively.

**Fig 3:**
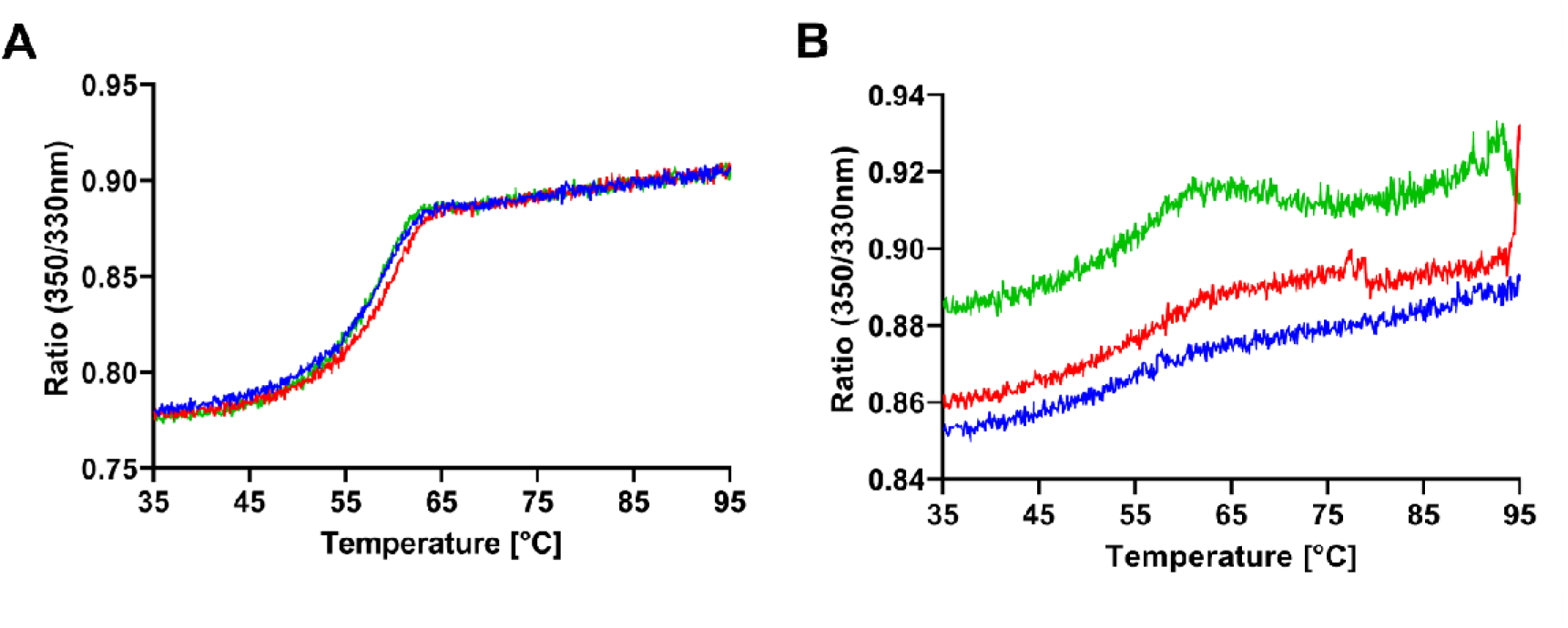
Differential scanning fluorimetry - Ratio of intrinsic fluorescence plotted as a function of temperature for refolded 4B7-Db (A) and 1245-Db (B). The inflection temperature of 4B7-Db was 59.07 ± 1°C, which is the average of three independent experiments, with each experiment run in triplicates. The variability in each of the triplicates in 1245-Db could be due to precipitation of protein reflected as a change in intensity, without a change in the overall profile.

It is speculated that the nature of amino acids, presence of additional tryptophan or alkaline pI, could be the reason behind the inability of 1245-Db, to refold in the same condition as 4B7-Db and 1269-Db, albeit it cannot be ascertained.

### 4B7-Db showed dimeric species in solution when refolded from inclusion bodies

The inclusion of a five amino acid linker, between the variable heavy (V_H_) and variable light (V_L_)chains, leads to the association of dimers, called diabodies (4B7-Db) to form a functional antibody fragment. Consistent with the design, existence of diabody in solution, which increased with incubation time, was demonstrated with a glutaraldehyde cross linking assay, visualized on Coomassie stained SDS-PAGE gel (Fig 4A)(8). However, in the case of 1245-Db, no time dependent increase in dimeric species was observed (Fig. 4B). This could be due to a small fraction of refolded species present in solution, forming a diabody in 1245-Db (Fig. 4B). PfCelTOS, is reported to exist as an obligate dimer in solution, was predominantly present as a dimeric fraction after 30 minutes of incubation (Fig. 4C).

**Fig 4:**
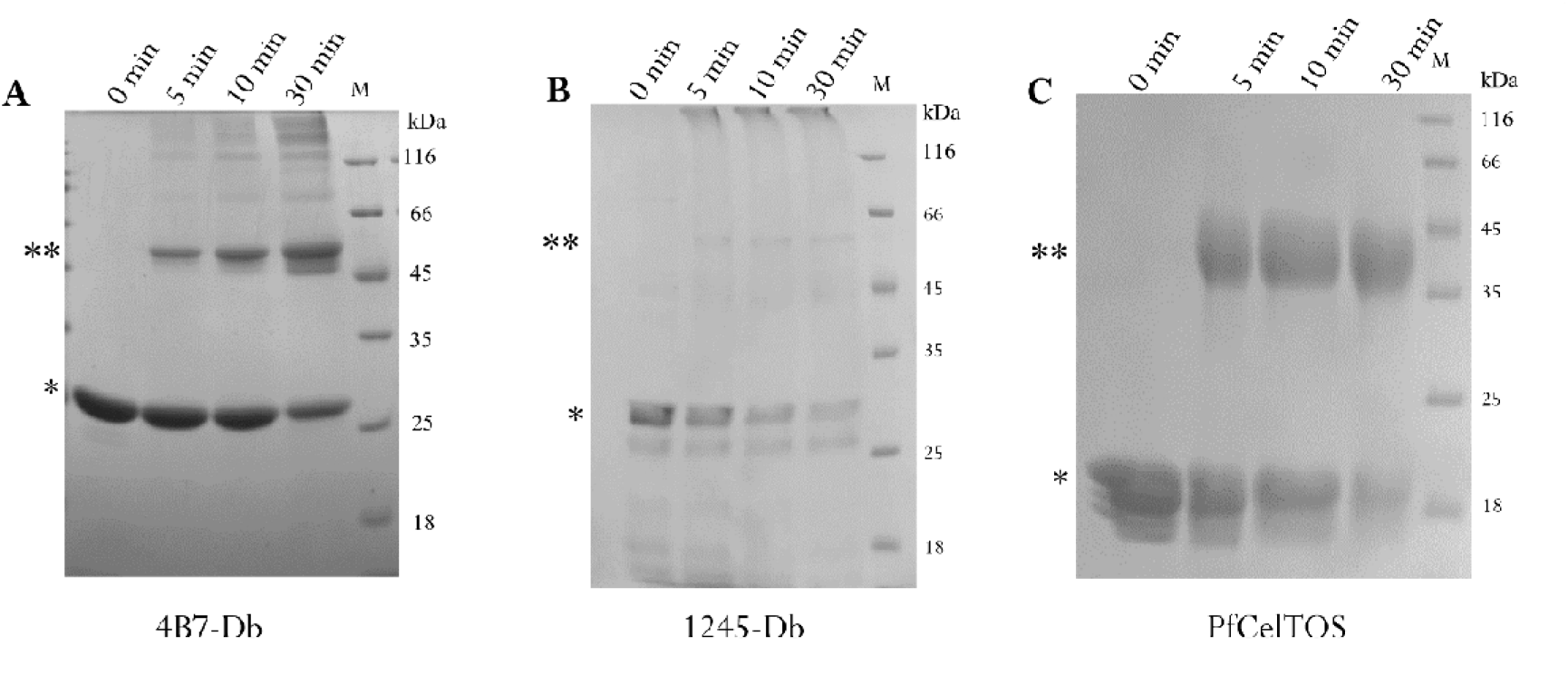
A & B - Glutaraldehyde cross linking of 4B7-Db and 1245-Db in solution, respectively. For 4B7-Db, the proportion of dimeric species (**) increases with incubation time, with a decrease in the proportion of monomeric species (*). A thin band is seen corresponding to the dimeric fraction in 1245-Db, the intensity of which does not increase with time. PfCelTOS is an obligate dimer in solution, which gets crosslinked at 5 minutes, used in this experiment as a positive control.

### Refolded 4B7-Db recognized surface expressed and purified Pfs25

4B7-Db exhibited a fluorescence signal when incubated with ookinetes, indicating its recognition of surface expressed Pfs25 (Fig 5). However, 1245-Db showed weak recognition of Pfs25 on ookinetes (Fig. 5). This result was reproduced from two independent batches of refolded 1245-Db. As 1245-Db exhibited a substandard DSF profile, without detectable diabody and recognized Pfs25 poorly on the surface of ookinetes, we did not use this antibody fragment for binding studies with purified MBP-Pfs25.

**Fig 5:**
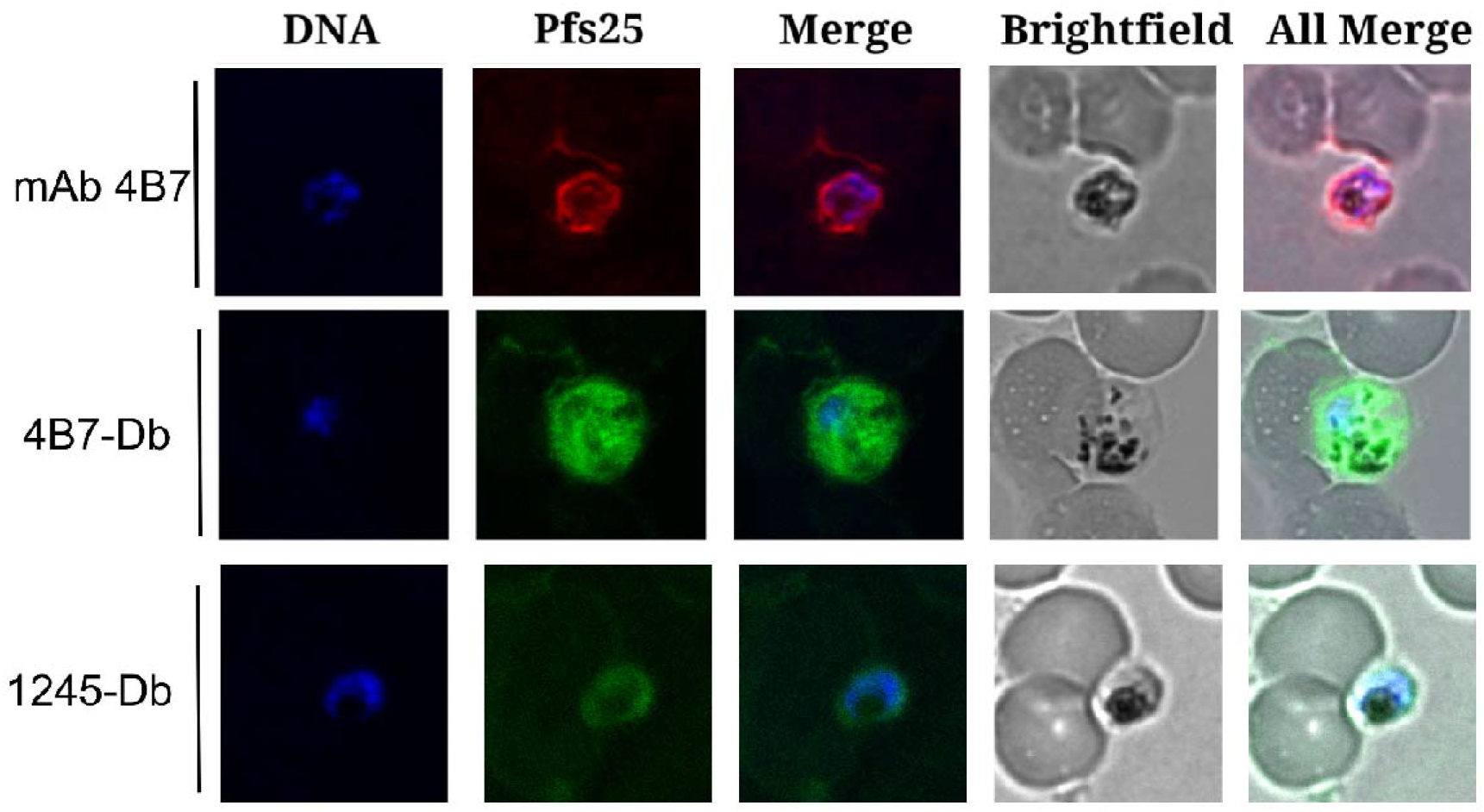
Immunofluorescence based Binding assays with refolded 4B7-Db and 1245-Db. Monoclonal antibody mAb 4B7 was used as a control along with 1245-Db as the test sample for IFA. Diabody 4B7-Db shown in the second panel was evaluated alongside 1269-Db and mAb 4B7 as shown in figure 9 of Jagannath et. al (19). A small fraction of 1245-Db diabody observed in glutaraldehyde cross-linking assay could give rise to the weak signal observed in IFA. Further, the background signal (green) is higher in 1245-Db panel, implying the specific signal to be lower than observed.

4B7 monoclonal antibody, recognizes a linear epitope on Pfs25 (15), due to which it can recognize denatured Pfs25, *in situ* on a western blot (19). In line with its parent antibody, 4B7-Db recognized denatured Pfs25, with a western blot based binding assay (Fig 6A). We had earlier shown that 1269-Db, recognizing a conformational epitope on Pfs25, failed to exhibit binding with denatured Pfs25(19). In addition to recognizing denatured epitope, 4B7-Db also showed binding to native monomeric MBP-Pfs25 (Fig. 6B)(19).

**Fig 6:**
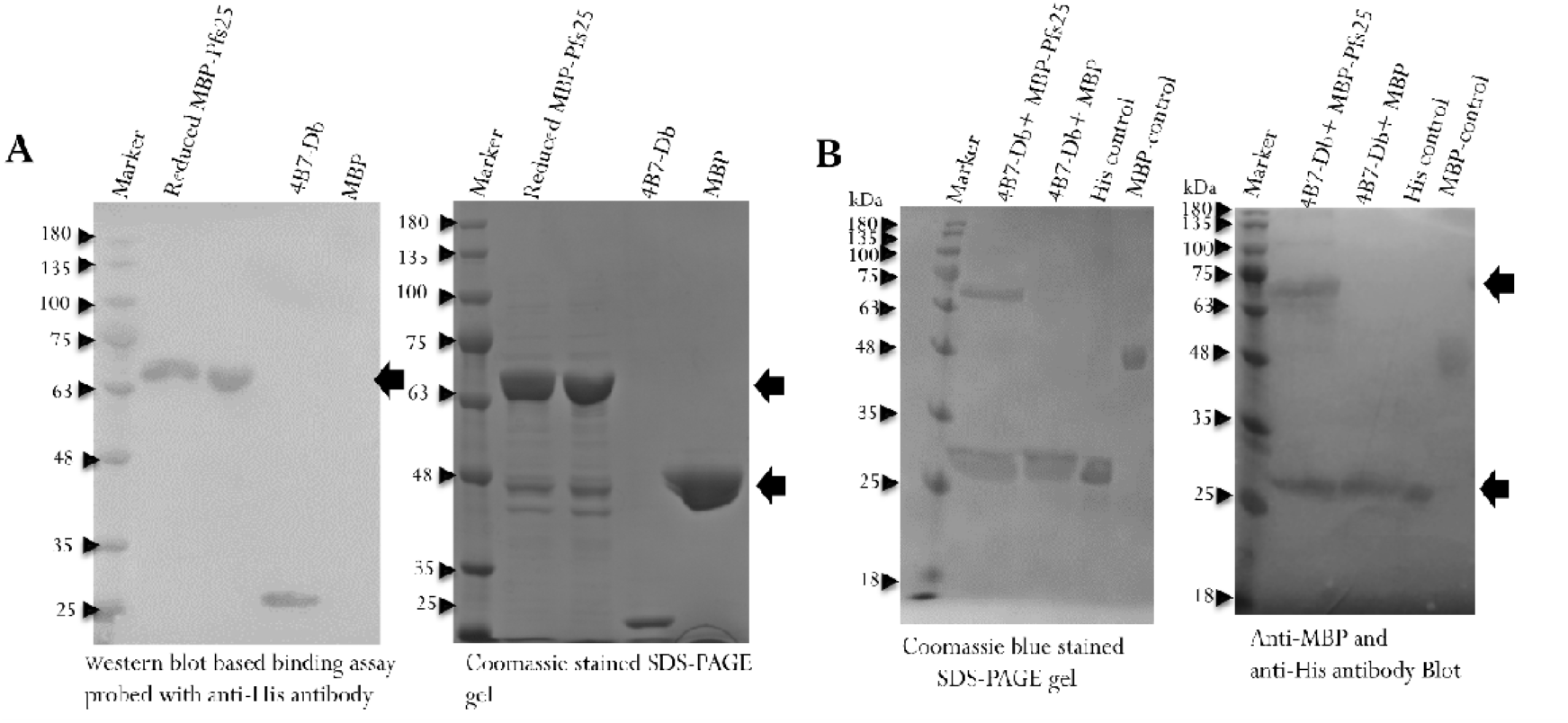
Binding of purified Pfs25 with 4B7-Db. A. Western blot based binding assay: purified MBP-Pfs25 (under reduced and non-reduced conditions) and MBP run on an SDS-PAGE gel (right panel) was transferred onto a blot and incubated with 4B7-Db. The histidine tagged 4B7-Db bound to Pfs25, was recognized by anti-His conjugate as described in methods. MBP was not recognized by 4B7-Db. B. Pulldown assay with native monomeric MBP Pfs25 protein. Pfs25 interacting with Histidine tagged 4B7-Db bound to Ni-NTA column was eluted, run on an SDS-PAGE gel (left panel) and probed with both anti-His and anti-MBP antibody in western blot. 4B7-Db bound to monomeric MBP-Pfs25 but not MBP, as seen in the blot and gel.

## Discussion

Mortality and morbidity due to malaria continues to be high in Sub-Saharan Africa, despite preventive and therapeutic interventions made available in the region (20). This was further exacerbated by the COVID19 pandemic (21). A multi-combinatorial approach with different preventive and therapeutic strategies, is needed for the elimination of malaria. Recently, a human monoclonal antibody, CIS43LS targeting circumsporozoite protein (CSP) of the parasite is in Phase 1 clinical trials for passive immunization (13). Parasite transmission is blocked inside the mosquitoes, when the antibodies generated after administration of transmission blocking vaccine candidates, are ingested by the mosquitoes during blood feeding (22). 4B7 and 1245 are transmission blocking antibodies, generated by immunization of mice with the *Plasmodium* protein, Pfs25 (17, 18). A combination of CIS43LS and a transmission blocking antibody, can protect individuals from the disease and reduce parasite load, consequently reducing the incidence of malaria. There is a need to design and engineer antibodies with desirable properties in preclinical and clinical phases, for them to be employed successfully in passive immunization (14). Antibody fragments retaining their antigenic specificities can be engineered to address the problems associated with therapeutic antibodies. Antibody fragments can be expressed in high levels in bacteria, the most sought expression system for its simplicity and low turnaround time (23). In this study, we designed two diabodies, with a five amino acid linker between V_H_ and V_L_ domains and refolded them using an optimized refolding condition, reported in our earlier study (19). Success in refolding antibody fragments from inclusion bodies is dependent on the refolding process and conditions. An undesirable environment for protein folding or a faulty design can increase the proportion of unfolded species. Methods which can assess the folded protein qualitatively, can be potentially used for preliminary characterization of purified proteins. We assessed the quality of refolded 4B7-Db and 1245-Db, using differential scanning fluorimetry (21, 22). As compared to isothermal denaturation analysis of proteins using fluorimetry and circular dichroism, DSF has many advantages including requirement of minimal quantity of protein for measurement and a faster turnaround time (24). DSF can be performed using a dye which binds to hydrophobic surface on protein (conventional DSF) or using intrinsic fluorescence of tryptophan residues (nanoDSF) (25). It has been used to identify buffer conditions in which the purified proteins are stable for biophysical studies including crystallization, and to qualitatively determine binding to small molecules, peptides and nucleic acids (25, 26, 27, 28). DSF has been reported to screen and identify refolding conditions in which the protein is folded optimally, reflected by a low initial fluorescence and sigmoidal unfolding transition of the melt profiles (29, 30). 4B7-Db showed a melting temperature of 59.07 ± 1°C determined using nanoDSF, with a low initial fluorescence and sigmoidal melt profile. In another report, the melting temperature of refolded scFvs, against the rabies virus glycoprotein was determined to be between 41 and 43°C (31). A correlation was observed between the melt profiles and functionality for 4B7-Db and 1245-Db (32, 33), indicating that DSF can be used for determining the quality of purified therapeutic antibody fragments. DSF, combined with mutagenesis, can be used to generate variants with specific applications for antibody fragments (34). Antibody fragment of 4B7, when expressed inside mosquitoes, blocked the transmission of *plasmodium* (35), suggesting that single chain antibodies can be used for transmission blocking immunization strategy, the aim of which is to reduce the parasite load in the population.

## Methods

### Cloning, expression, and refolding of 4B7- and 1245-Db

The gene sequences of V_H_ and V_L_ from the monoclonal antibody, 4B7 (GenBank: ADU55810.1) and 1245 (PDB ID: 6B0G), along with a G_4_S linker and a C-terminal hexa-histidine tag was codon optimized for expression in *E. coli* and subcloned into pET21a vector. Transformants of BL21(DE3) were grown to an OD of 0.8 in Luria Bertani medium with 100 ug/ml ampicillin and induced with 1 mM IPTG for 5 hours at 37°C. Inclusion body was prepared, solubilized and refolded with 50 mM tris pH 6.0, 9.6 mM NaCl, 0.4 mM KCl, 2.0 Mm MgCl2, 2.0 mM CaCl2, 0.5 M arginine, 0.05% polyethylene glycol 3350, 1 mM GSH and 0.1 mM GSSH for 24 hours, as described before (19). The purified proteins were visualized on a Coomassie stained SDS-PAGE gel and used for further analysis. MBP-Pfs25 was expressed and purified as reported earlier (19).

### Differential scanning fluorimetry

Protein melting profiles using 5 μM of 4B7- or 1245-Db were measured with Tycho NT.6 (NanoTemper® Technologies GmbH). Fluorescence emission intensities was measured at I_350_ and I_330_ in triplicates and its ratio plotted as a function of temperature, from 35 °C to 95 °C, with a 30 °C/min increase rate (NanoTemper® Technologies GmbH). The Inflection temperature (T_i_) was computed by the software based on melt curve analysis (31).

### Binding assays

For western blot-based binding and pull-down assays, MBP-Pfs25 was expressed in the periplasmic fraction and monomeric fraction purified with size exclusion chromatography, as described elsewhere (19). Western blot based binding assay and pulldown assays were performed as described before (19). Briefly, for western blot based binding assay, the PVDF transferred MBP-Pfs25 was probed with purified 4B7-Db and subsequently with anti-His antibody (1:3000), followed by Horse radish peroxidase -conjugated IgG (1:6000). The blot was developed with 3,3’,5,5’-Tetramethylbenzidine (TMB).

For pulldown assays, MBP-Pfs25 was incubated with histidine-tagged 4B7-Db, passed through Ni-NTA spin column (Takara Clontech), washed and column eluted with 500 mM Imidazole. The elutes were visualised with a Coomassie stained SDS-PAGE gel (12%). The elutes were also transferred onto a PVDF membrane and probed with anti-MBP + anti-His antibody. MBP was used as a negative control in both the binding assays to detect non-specific binding of antibody fragments (19).

### Immunofluorescence assay

Culture of *Plasmodium falciparum* (MR4) was grown till the zygote ookinete stage. Thin blood smears containing zygotes and ookinetes were prepared on glass slides and fixed with chilled methanol. Blocking and permeabilization were carried out simultaneously by incubating with 5% BSA and 0.2% saponin in 1X PBS for 60 mins at 25 °C. 5 μ g of antibody fragments or 4B7 (BEI) were added to the washed slides and incubated with (in 5% BSA and PBS) at 4 °C for 16 hours, and subsequently probed with alexa fluor 488-conjugated anti-His or anti-mouse IgG (H + L) Cross-Adsorbed Secondary Antibody Alexa Fluor 568 (ThermoFisher), respectively. Slides were mounted with Prolong Gold antifade along with DAPI for staining DNA (Thermo Fischer) and examined using a fluorescence microscope (19, 36).

### Ethics approval and consent to participate

Not applicable

### Consent for publication

Not applicable

## Acknowledgements

The following reagents were obtained through BEI Resources, NIAID,NIH: Monoclonal Anti-Plasmodium falciparum 25-kDa Gamete Surface Protein (Pfs25), Clone 4B7 (produced in vitro), MRA-28, contributed by Louis H. Miller and Allan Saul. *Plasmodium falciparum*, Strain NF54 (Patient Line E), MRA-1000, contributed by Megan G. Dowler. The authors are thankful to Dr Suresh Subramani and Dr Ethan Bier, University of California, San Diego for the valuable scientific discussions.

## Author’s contributions

Ashwathi Valiyaparambil optimised the process of refolding and conducted cross-linking experiments. Deepak K Jagannath refolded the 4B7-Db, purified MBP-Pfs25, performed differential scanning fluorimetry, purified MBP-Pfs25 and pull-down assays. Vysakh K Viswanath refolded 1245-Db. Sabyasachi Pradhan performed immunofluorescence assay for 4B7. Naveen Kumar revived and maintained the parasite culture and performed immunofluorescence assay for 1245-Db. Jay Prakash Shukla did the confocal imaging of the IFA slides. Anirudha Lakshminarasimhan designed the study and wrote the paper.

## Availability of data and materials

All the relevant data used to support the findings of this study are included within the article.

